# The outcome of neutrophil-T cell contact differs depending on activation status of both cell types

**DOI:** 10.1101/2020.11.05.370171

**Authors:** Danielle Minns, Katie J Smith, Gareth Hardisty, Adriano Rossi, Emily Gwyer Findlay

## Abstract

Neutrophils and T cells exist in close proximity in lymph nodes and inflamed tissues during health and disease. They are able to form stable interactions, with profound effects on the phenotype and function of the T cells. However, the outcome of these effects are frequently contradictory; in some systems neutrophils suppress T cell proliferation, in others they are activatory or present antigen directly. Published protocols modelling these interactions *in vitro* do not reflect the full range of interactions found *in vivo*; they do not examine how activated and naïve T cells differentially respond to neutrophils, or whether de-granulating or resting neutrophils induce different outcomes. Here, we established a culture protocol to ask these questions with human T cells and autologous neutrophils. We find that resting neutrophils suppress T cell proliferation, activation and cytokine production but that de-granulating neutrophils do not, and neutrophil released intracellular contents are pro-activatory. Strikingly, we also demonstrate that T cells early in the activation process are susceptible to suppression by neutrophils, while later-stage T cells are not, and naïve T cells do not respond at all. Our protocol therefore allows nuanced analysis of the outcome of interaction of these cells and may explain contradictory results observed previously.

## Introduction

Neutrophils and T cells co-exist in the lymph nodes and tissues during health and disease^1–4^. Stable interactions between the two have been observed^5^, with profound effects on the phenotype and function of the T cells. However, the data is confusing; in some studies neutrophils can activate T cells, present antigen directly and enhance proliferation^6–10^ while in others neutrophils suppress proliferation and induce apoptosis^5,11–13^.

It is critically important to understand the mechanisms behind these interactions and how they may alter T cell function during disease, and in particular to do so using human cells. Furthermore, deciphering confusing *in vivo* results by performing depletion of neutrophils is difficult as this leads to an increase in T cell-stimulatory cytokine production^14^. As a result, *in vitro* co-culture systems are essential for unpicking mechanisms by which neutrophils influence T cell behaviour.

Previous work using such systems has cultured naïve T cells with untouched, freshly isolated peripheral blood neutrophils, usually in the presence of high-dose T cell activation agents. These cultures mimic the interaction of resting neutrophils with early-activating T cells undergoing antigen presentation by dendritic cells in the lymph nodes.

These cultures do not however model other interactions that occur *in vivo*: 1) naïve cell-cell contact in the blood, 2) interaction of T cells with activated neutrophils or their released contents during the c.24 hours T cells are receiving signals in lymph nodes^4,15–17^ or 3) contact in inflamed tissues or tumours, at which sites the vast majority of T cells present have previously been activated in the lymph nodes and are not naive.

In addition, neutrophils moving into lymph nodes and tissues during disease are often activated or releasing their contents by de-granulation or NETosis^18^. To our knowledge, no *in vitro* paper has modelled the interaction of human T cells with autologous primed neutrophils or their released contents. Here, we describe a novel protocol for these co-cultures and assess the impact of differing neutrophils on T cell phenotype and function.

Our protocol is shown in Fig. 1A. T cells and autologous neutrophils are isolated from peripheral blood of healthy donors within 30 minutes of blood draw. To model interaction of naïve T cells and resting neutrophils (such as occur in the blood), cells are co-cultured without stimulation. To model the interaction of early-activating T cells and neutrophils in the lymph node, in the presence of dendritic cells bearing foreign antigen, cells are co-cultured in the presence of a low dose of αCD3/αCD28 activator. To model interaction of late-activating T cells with neutrophils in the inflamed tissue, T cells are first given αCD3/αCD28 stimulation for 24 hours in the form of Dynabeads. The beads are then removed with magnets, and fresh autologous neutrophils added. This avoids the use of Dynabeads in the same well as neutrophils, which can inhibit Dynabead action and give the false appearance of suppression^19^; however, use of Dynabeads rather than soluble activators for activation in the absence of neutrophils allows complete removal with magnets before neutrophils are added on day 1.

**Figure 1:**
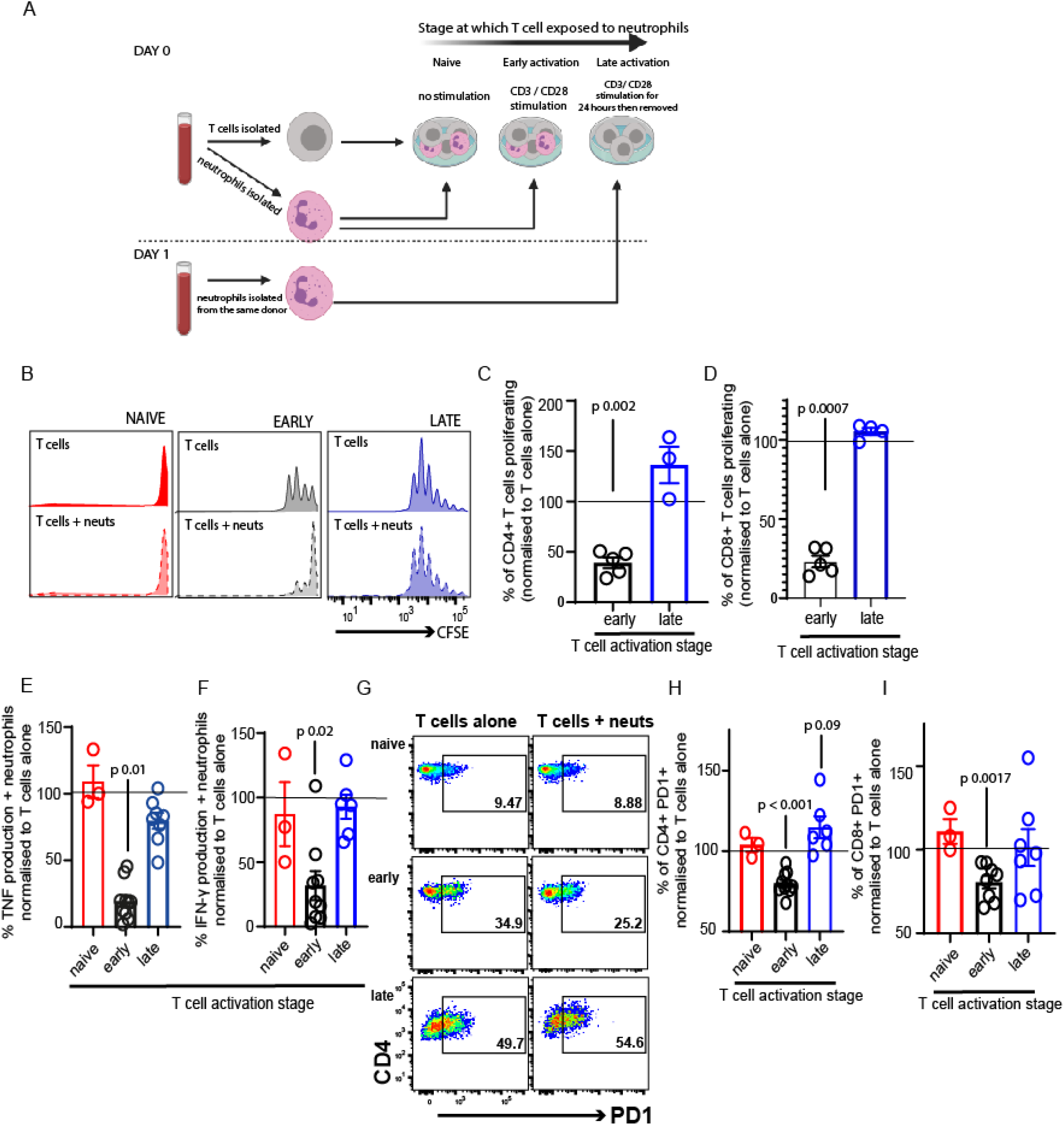
Neutrophil contact differentially affects early- and late-activating T cells. (A) Schematic of culture system. Rapid isolation of peripheral blood neutrophils by negative magnetic selection is performed before autologous T cells are isolated. Cells are co-cultured at a 5:1 neutrophil: T cell ratio either without any stimulation (‘naive’) or in the presence of CD3 / CD28 activation cocktail (‘early’). T cells are also cultured alone in the presence of CD3/ CD28 coated Dynabeads for 24 hours. The next day Dynabeads are removed and fresh autologous neutrophils isolated within 30 minutes of blood draw. These are co-cultured without any further stimulation (‘late’). (B-D) Proliferation of T cells is assessed by measuring CFSE dye dilution following 72 hours’ co-culture. (E-F) Culture supernatant is collected following 24 hours’ co-culture to assess cytokine production by ELISA. (G-I) Activation is assessed following 24 hours’ co-culture by measuring PD1 expression by flow cytometry. Data shown is individual donors, bars denote standard error. N = 3-8. Statistical tests used: C,D, E,F, H, I: paired two-tailed T tests between neutrophil-exposed and control wells, all tests done on raw data before conversion to %.

We assess the impact of neutrophils which are resting or primed, as well as the intracellular contents of the neutrophils^20^, on T cell phenotype.

Our protocol therefore allows analysis of the many ways T cells and neutrophils can interact during inflammatory disease. It more closely models *in vivo* situations than any other currently used. Using this protocol, we asked two questions: 1. Do naïve, early-activating and late-activating T cells respond differently to neutrophil contact? And 2. How do T cells respond to resting or primed neutrophils or their contents?

## Results and discussion

### Early- and late-activating T cells respond differently to neutrophil contact

We firstly examined the impact of neutrophil exposure on T cell proliferation after 72 hours’ co-culture. Naïve T cells did not proliferate either in the presence or absence of neutrophils (Fig. 1B). As previously shown, neutrophils suppressed proliferation of early-activating CD4^+^ and CD8^+^ T cells (Fig. 1B-D). However, this was not the case with late-activating T cells. If the T cells had received stimulation for 24 hours before neutrophil addition, neutrophils had no impact on their proliferation over the subsequent 72 hours, and even slightly increased the proliferation of CD8^+^ T cells (Fig. 1B-D).

Culture supernatants were collected to assess production of inflammatory cytokines following 24 hours’ co-culture. T cell production of TNF (Fig. 1E) and IFN-γ (Fig. 1F) was strikingly reduced by exposure to neutrophils, if it occurred during the early activation process. However, when late-activation stage T cells were incubated with neutrophils, a reduction in further cytokine production did not occur. The very low concentration of cytokines produced by naïve T cells was not altered by neutrophil exposure.

Next, we examined the expression of PD1, an early marker of T cell activation^21,22^. Naïve T cells expressed low levels of PD1 after 24 hours in culture, and this was unchanged by neutrophil exposure (Fig. 1G,H,I). Neutrophils suppressed activation of T cells in the early-activating cultures, with the proportion expressing PD1 reducing significantly (Fig. 1G,H,I). In contrast, neutrophil presence increased the frequency of PD1 expression on late-activating CD4+ T cells (Fig. 1H), and did not suppress expression on CD8+ late-activating cells (Fig. 1I).

Together, this suggests that early lymph node-stage T cells encountering neutrophils are highly susceptible to suppression of proliferation, activation and cytokine production. In contrast, late tissue-stage T cells are not suppressed by neutrophil contact. In contrast, proliferation and activation of T cells at the ‘tissue-stage’ can be enhanced by contact with resting neutrophils.

### Primed and resting neutrophils induce opposite responses in T cells

Our second question was whether resting or primed neutrophils, or their contents, differentially affected T cells. Strikingly, while resting neutrophils suppressed early T cell activation (Fig. 1H,I and Fig. 2B,C), primed neutrophils did not. Neutrophil contents also increased PD1 expression on CD8^+^ cells (Fig, 2C), and induced significantly different outcomes to resting neutrophils in both T cell subsets (Fig. 2B,C).

**Figure 2:**
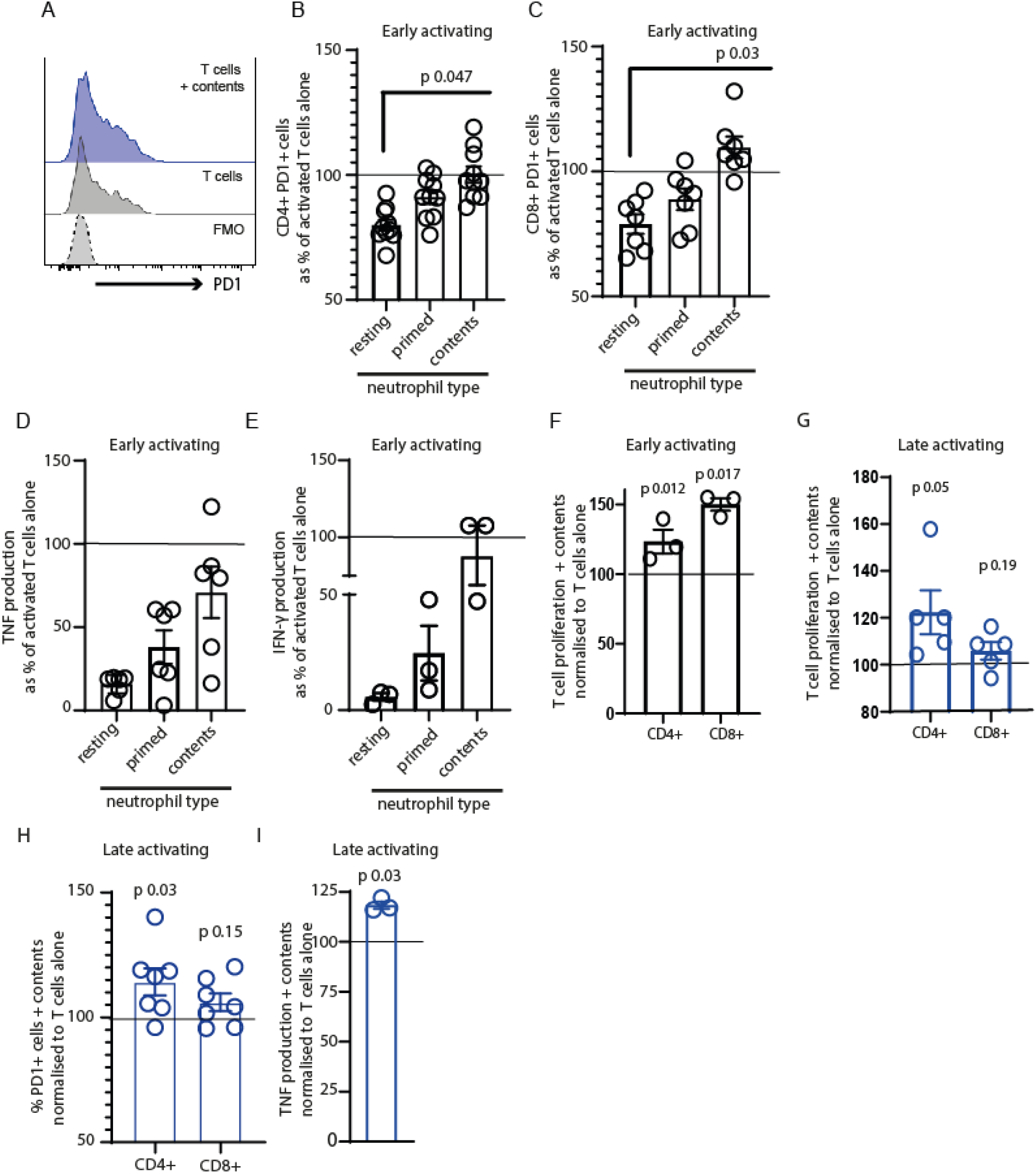
Primed and resting neutrophils, and their released contents, differentially alter T cell phenotype. T cells were plated with autologous neutrophils In a 5:1 neut:T cell ratio. Neutrophils were either resting (untouched), primed (10uM cytochalasin B, lOOnM fMLF) or their contents released by 5 rapid freeze-thaw cycles. After 24 hours’co-culture activation was assessed by measuring PD1 expression by flow cytometry (A-C). (D-E) cell culture supernatants were collected and cytokines measured by ELISA. (F-G) Teel Is were labelled with CFSE and their proliferation measured by Its dilution following 72 hours’ co-culture. Tcells were activated for 24 hours then the activation beads removed and fresh autologous neutrophils added. Activation (H and cytokine production (I) were measured 24 hours later. Data shown are Individual donors with bars representing standard error of the mean. N= 4-10.Tests used: B, C: one-way ANOVA with Dunnett’s post test comparing all to resting neutrophils; F, G, H, I,: paired two-tailed t tests comparing to activated T cells alone, tests performed on raw data.

Primed neutrophils or their contents did not suppress T cell TNF (Fig. 2D) or IFN-γ (Fig. 2E) production as much as resting cells, with contents inducing almost no suppression at all, suggesting that this is dependent on cell-cell contact.

We were interested in the non-suppressive nature of neutrophil contents on early-activating T cells and investigated their impacts in more detail. Surprisingly, neutrophil contents enhanced the proliferation of both early-activating (Fig. 2F) and late-activating (Fig. 2G) T cells. Late-activating T cells also increased their PD1 expression and proliferation following contact with contents (Fig. 2H). In addition, the production of TNF (Fig. 2I) was enhanced.

We conclude that neutrophils have no impact on naïve T cells, as might be expected for cells that meet so frequently in the blood. We show that if resting neutrophils are present early in the T cell activation process, they strongly suppress activation, proliferation and cytokine production. However, later on in this process they do not suppress, and can in fact enhance activation of T cells.

However, if the neutrophils present at the early stage are primed, or have released their intracellular contents, they are not suppressive. This suggests a check in the development of an adaptive response, in which neutrophil activatory signals must have been received to promote an inflammatory response but the default is suppression.

Finally, we demonstrate that the release of neutrophils’ contents, which can occur during NETosis, degranulation and death, enhances proliferation of T cells and, in particular, is strongly pro-stimulatory to late-stage activating T cells. This is a model of encounter of interactions in the inflamed tissue.

To our knowledge this is the first demonstration that neutrophil contents have opposite outcomes on T cell phenotype to resting cells, and that T cell activation stage differentially affects outcome. Our work confirms the need for culture models which recapitulate each stage of inflammatory disease with accuracy, and may explain why *in vitro* systems used previously do not always explain observations made *in vivo*.

## Authorship contributions

DM, KJS, and EGF performed experiments; DM, GH, AR and EGF analysed data and wrote the manuscript.

## Acknowledgments

We thank the QMRI flow cytometry team (Shonna Johnston, Will Ramsay and Mari George) for their help and advice.

## Conflict of interest statement

The authors declare no conflicts of interest.

## Methods

### Healthy Human Donors

Peripheral venous blood was collected from healthy adult volunteers under ethical agreement code AMREC 20-HV-069, which included informed written consent. Blood was collected into sodium citrate and was processed immediately or within 30 mins of blood draw.

### T Cell and Neutrophil EasySep Isolation

EasySep separation kits (StemCell Technologies, T Cells: #19661, Neutrophils: #19257) were used to isolate CD3^+^ T cells or neutrophils directly from human whole blood, as per the manufacturer’s guidelines.

### Neutrophil Treatments

We assessed the impact of resting and primed neutrophils, as well as neutrophil contents. Neutrophils were resuspended in 1X PBS at a concentration of 7.5 million/mL. Resting neutrophils were untreated and cultured with T cells immediately. Primed neutrophils were obtained by treating the cells for 25 mins with 10 μM cytochalasin B (Merck, #C6762) and 100 nM N-Formylmethionyl-leucyl-phenylalanine (fMLF) (Merck, #F3506). Cells were then washed well before co-culture. Neutrophil contents were obtained by freeze-thawing the cells 5 times on dry ice, followed by high speed spin to remove cell membranes, as described by Miles et al^20^.

### Cell Culture

T cells were co-cultured with neutrophils at a 5:1 N:T ratio in round-bottom 96 well plates, in complete medium (RPMI, 10% fetal calf serum, 10 units/mL penicillin, 10 μg/mL streptomycin and 2 mM L-glutamine, all supplied by Gibco, ThermoFisher UK). To model interaction of naïve T cells and resting neutrophils, cells were co-cultured in the absence of any stimulation. To model the interaction of early-activating T cells and neutrophils in the lymph node, cells were co-cultured in the presence of αCD3/αCD28 activator (1 μL ImmunoCult activator to 1×10^5^ cells, StemCell, #10971), which remained in the well throughout culture. To model interaction of late-activating T cells with neutrophils in the inflamed tissue, T cells were first given αCD3/αCD28 stimulation for 24 h in the form of CD3/CD28-coated Dynabeads (1 bead:100 cells; Gibco, #11161D). The beads were then removed with a magnet, the same donor bled again, and fresh autologous neutrophils added as above in fresh medium.

### Proliferation Assay

In order to assess proliferation, T cells were resuspended in 1X PBS and stained with CFSE (Invitrogen, #C34554, working concentration: 5 μM). T cells were incubated at 37°C for 20 mins and then washed twice with an excess of media. Proliferation analysis by dye dilution was performed by flow cytometry following 3 days co-culture.

### Flow Cytometry

Cells were stained for surface markers for 30 mins at 4°C, protected from light. DAPI (Invitrogen, #D1306, working concentration: 1 μg/mL) was added prior to running to assess viability. Samples were analysed using a Fortessa LSR II (BD Biosciences) and FlowJo software (Treestar).

### Antibodies

CD4-PE/Cy7 (clone A161A1, Biolegend, #357410, lot B225074); CD8-AF647 (clone HIT8a, Biolegend, #300918, lot B235677); PD1-PE (clone EH12.2A7, Biolegend, #329905, lot B252642).

### ELISAs

The concentration of TNFα (R&D DuoSet ELISA, #DY210) and IFNγ (R&D DuoSet ELISA, #DY285) in cell culture supernatants was determined by ELISA, as per the manufacturer’s guidelines.

### Statistics

All data shown are expressed as individual data points with line at mean +/- standard error. Statistical analysis was performed using GraphPad Prism software. Two groups were compared with two-way paired Student’s t-tests. Multiple groups were compared using a one-way analysis of variance (ANOVA) test with a Dunnett post-test. A minimum of three donors were used over at least 2 experiments. Details of sample sizes and statistical analyses performed are included in all figure legends.

### Data sharing statement

All data are available upon request to the corresponding author at

## Notes

### Competing Interest Statement

The authors have declared no competing interest.

